# Classification and monomer-by-monomer annotation of suprachromosomal family 1 alpha satellite higher-order repeats in hg38 human genome assembly

**DOI:** 10.1101/408674

**Authors:** L. Uralsky, V.A. Shepelev, A.A. Alexandrov, Y.B. Yurov, E.I. Rogaev, I.A. Alexandrov

## Abstract

In the latest hg38 human genome assembly, centromeric gaps has been filled in by alpha satellite (AS) reference models (RMs) which are statistical representations of homogeneous higher-order repeat (HOR) arrays that make up the bulk of the centromeric regions. We studied these models to compose an atlas of human HORs where each monomer of a HOR could be characterized and represented by a number of its polymorphic sequence variants. We further used these data and HMMER sequence analysis platform to annotate AS HORs in the assembly. This led to discovery and annotation of a new type of low copy number highly divergent HORs which were not represented by RMs. The annotation can be viewed as UCSC Genome Browser custom track (the HOR-track) and used together with our previous annotation of AS SFs in the same assembly where each AS monomer can be viewed in its genomic context together with its classification into one of the 5 major SFs (the SF-track). To catalog the diversity of AS HORs in the human genome we introduced a new naming system. Each HOR received a name which showed its SF, chromosomal location and index number. Here we present the first installment of the HOR-track covering only the 17 HORs that belong to SF1 which forms live functional centromeres in chromosomes 1, 3, 5, 6, 7, 10, 12, 16 and 19 and also a large number of minor dead HOR domains, both homogeneous (pseudo) and divergent (relic). The 4 newly discovered divergent SF1 HORs have provided the missing links in SF1 early evolution and substantiated its partition into 2 generations, archaic and modern, which we reported earlier. Additionally, we demonstrated that monomer-by-monomer HOR annotation was useful for mapping and quantification of various structural variants of AS HORs which would be important for studies of inter-individual polymorphism of AS including centromeric functional epialleles.

---

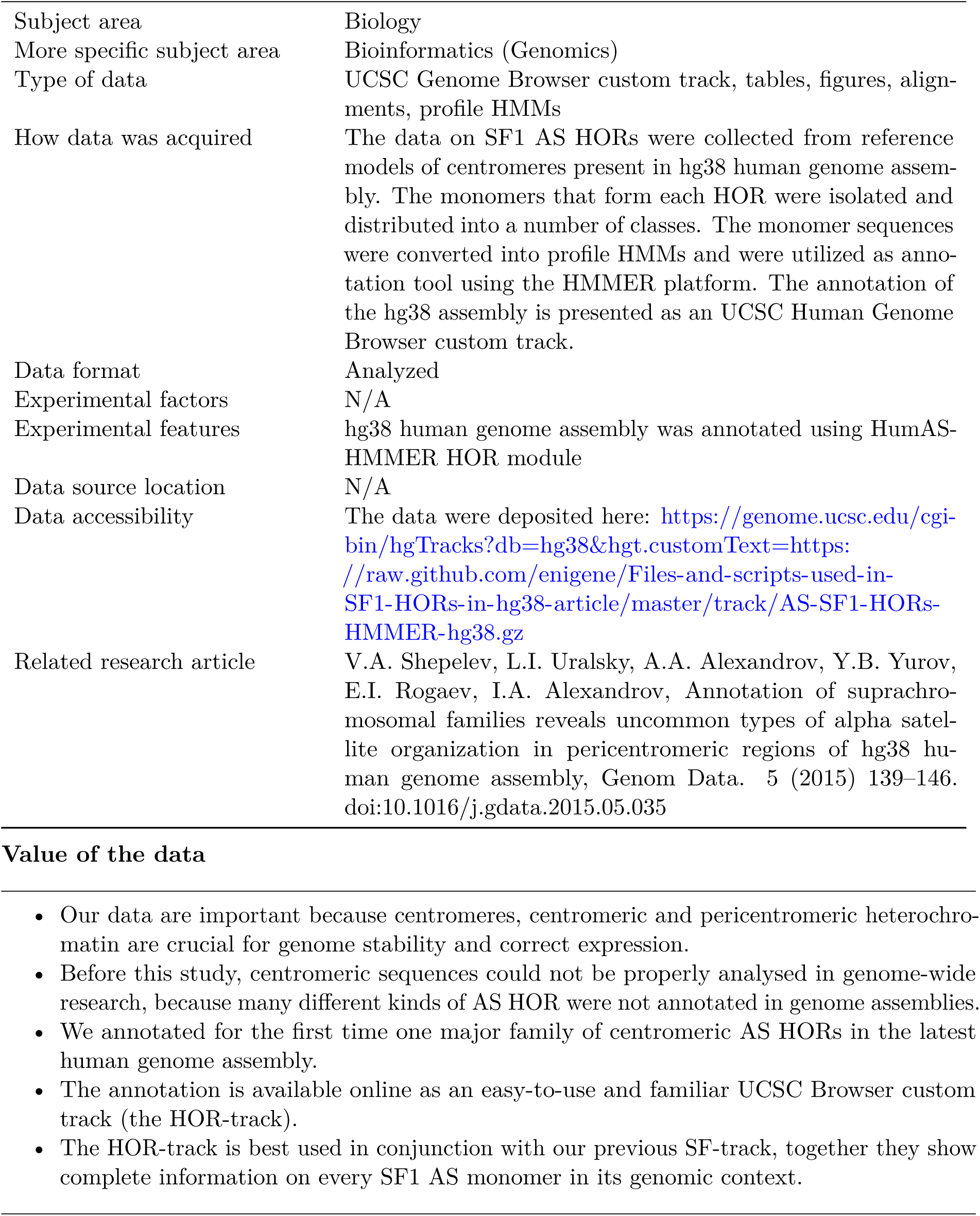

|  |  |
| --- | --- |
| Subject area | Biology |
| More specific subject area | Bioinformatics (Genomics) |
| Type of data | UCSC Genome Browser custom track, tables, figures, alignments, profile HMMs |
| How data was acquired | The data on SF1 AS HORs were collected from reference models of centromeres present in hg38 human genome assembly. The monomers that form each HOR were isolated and distributed into a number of classes. The monomer sequences were converted into profile HMMs and were utilized as annotation tool using the HMMER platform. The annotation of the hg38 assembly is presented as an UCSC Human Genome Browser custom track. |
| Data format | Analyzed |
| Experimental factors | N/A |
| Experimental features | hg38 human genome assembly was annotated using HumAS-HMMER HOR module |
| Data source location | N/A |
| Data accessibility | The data were deposited here: https://genome.ucsc.edu/cgi-bin/hgTracks?db=hg38&hgt.customText=https://raw.githubusercontent.com/enigene/Files-and-scripts-used-in-SF1-HORs-in-hg38-article/master/track/AS-SF1-HORs-HMMER-hg38.gz |
| Related research article | V.A. Shepelev, L.I. Uralsky, A.A. Alexandrov, Y.B. Yurov, E.I. Rogaev, I.A. Alexandrov, Annotation of suprachromosomal families reveals uncommon types of alpha satellite organization in pericentromeric regions of hg38 human genome assembly, Genom Data. 5 (2015) 139–146. doi:10.1016/j.gdata.2015.05.035 |

## 1. Data

This data article presents two kinds of material. Firstly, we report an exhaustive catalog of SF1 HORs in hg38 human genome assembly (17 HORs altogether). Each monomer is annotated to show its SF-specific monomeric class (176 kinds of monomers altogether). As each HOR is represented in the genome by a number of structural polymorphic variants, a separate catalog of such variants is provided for each HOR as represented in the corresponding RM. These data are collected in Table 1 and Fig. S1 and S2. Secondly, we annotated the 176 kinds of monomers identified through the entire hg38 assembly. These data are presented as UCSC Human Genome Browser custom track (the HOR-track). Direct link to deposited data: https://genome.ucsc.edu/cgi-bin/hgTracks?db=hg38-hgt.customText=https://raw.github.com/enigene/Files-and-scripts-used-in-SF1-HORs-in-hg38-article/master/track/AS-SF1-HORs-HMMER-hg38.gz

## 2. Experimental Design, Materials, and Methods

### 2.1. SF1 in morpho-functional classification of AS used in this work

AS is a satellite DNA that forms the centromeres of human chromosomes. It consists of tandem arrays of ~171 bp AS monomers. Classification of AS monomers has been recently summarized in [1]. There are 5 major AS SFs, of which the new SFs (1-3) form active (live) centromeres of all human autosomes and the X chromosome. The old SFs 4+ and 5 unite the sequences of the dead centromeres of our ancestors that lost centromeric function, but the remnants of which remained in the genome. SF4+ is an umbrella group which unites a large number of old and ancient SFs, such as SF4 proper, SF6, SF7 and more. The classification of SFs within SF4+ group is yet to be finalized, so for now they do not have formal names and are color-coded and can be described as colored pericentromeric layers [2]. SF5 is the immediate ancestor of the new SFs. It consists of the two monomeric classes R1 and R2 which represent two progenitor types (B and A, respectively) to which all monomeric classes of the new families including the two classes that form SF1 (J1 and J2) belong. Namely, R1 is the immediate ancestor of J2 and R2 of J1. SF1 is a major new family which forms live centromeres of chromosomes 1, 3, 5, 6, 7, 10, 12, 16 and 19 [3]. It consists of chromosome-specific HORs which include 2-18 monomers where J1 and J2 monomers usually appear in perfect or near perfect alternation and form internal J1J2 dimers. HORs are nearly identical units made by a number of somewhat different monomers (or dimers, trimers, etc.). As a rule, each chromosome has a tandem high copy number HOR of unique length and sequence (a chromosome-specific HOR) which forms the single live centromere in a given chromosome. However, some non-homologous pairs of chromosomes share almost identical or very similar live HORs (the so-called “paired domains” 13/21 and 14/22 and “triple domain” 1/5/19) [3]. Similarity between HOR copies is higher than between any repeated units within them (e.g. J1J2 dimers). Divergence between J1 and J2 classes is about 25% and within the classes (e.g. between J1J2 dimers within a HOR or from different HORs) may be about 12%. Divergence between copies of the same HOR is typically 1-2% [3].

Besides live homogenous centromeres, SF1 is found in different classes of dead (non-functional) arrays of which some are divergent (“dead relic centromeres”, first reported in this paper, see section 2.6 below) and some are homogenous (“dead pseudocentromeres” [1]). Live HOR arrays organize the kinetochore in most individuals and they are usually the largest HOR arrays in a given chromosome. However, in some individuals (or rather in some individual chromosomes), a smaller pseudocentromeric HOR array may assume the role of kinetochore organizer instead of the main array and form a centromeric epiallele [4, 5]. We propose that such occasionally functional HORs may be called “half dead” arrays as opposed to the “dead” ones that are never functional. Then, there are two slightly different possibilities: (1) half dead centromeres which have been live centromeres once, but have surrendered the status of the main centromere to a more efficient competitor and have retained only occasional activity; and (2) half dead pseudocentromeres which are the HORs that have never been live centromeres, but are recent amplifications of some dead AS sequences which occasionally assume centromeric activity. Note that divergent HORs are not known ever to be functional. So the dead relic HORs are probably dead for good unless some segments of them re-amplify and restore homogeneity which could make them eligible for a new status. For concise summary of AS organisation in human genome and in hg38 assembly see our recent paper [1] and references therein. The SF-track with annotation of AS monomers [1] is available at: https://genome.ucsc.edu/cgi-bin/hgTracks?db=hg38&hgt.customText=https://raw.github.com/enigene/AS-tracks/master/GRCh38-GCA_000001405.15/human-GRC-hg38-M1SFsv2.2.bed.gz. AS terminology we use is also summarized in [1].

**Table 1.** List of SF1 AS HORs in hg38 human genome assembly.

| # | Chrom. | New HOR name | Old HOR name | HOR length (mon) | RM or sample contig<sup>1</sup> | Genomic size (kb)<sup>2</sup> | Homogeneous/Divergent | Age | Status<sup>3</sup> | CENP-A reads<sup>4</sup> | Ref. |
| --- | --- | --- | --- | --- | --- | --- | --- | --- | --- | --- | --- |
| 1 | 1,5,19 | S1C1/5/19H1L<sup>5</sup> | D1Z7/<br>D5Z2/<br>D19Z3 | 6 | GJ212201.1 | 2282 | homogeneous | modern | live | 43,226 | [6] |
| 2 | 3 | S1C3H1L | D3Z1 | 17 | GJ211871.1 | 2102 | homogeneous | modern/<br>archaic | live | 31,967 | [7] |
| 3 | 3 | S1C3H2<sup>5</sup> | D3-2 | 10 | GJ211866.1 | 461 | homogeneous | archaic | pseudo | 484 | [1] |
| 4 | 3 | S1C3H3d | - | 5 | ABBA01004655.1 | 217 | divergent | archaic | relic | - | this paper |
| 5 | 3,6,7,8,<br>10,12,20 | S1CMH1d | - | 4 | ABBA01004652.1 | 251 | divergent | archaic | relic | - | this paper |
| 6 | 5p | S1C5pH2 | - | 16 | GJ211887.1 | 143 | homogeneous | modern | pseudo | 451 | RM |
| 7 | 6 | S1C6H1L | D6Z1 | 18 | GJ211907.1 | 1276 | homogeneous | archaic | live | 32,711 | [8] |
| 8 | 7 | S1C7H1L | D7Z1 | 6 | GJ211908.1 | 2659 | homogeneous | modern | live | 32,278 | [9] |
| 9 | 10 | S1C10H1L | D10Z1 | 8 | GJ211932.1 | 1561 | homogeneous | modern | live | 30,214 | [10] |
| 10 | 10 | S1C10H1-B | - | 14 | GJ211933.1 | 48 | homogeneous | modern | pseudo | 1,028 | RM |
| 11 | 10 | S1C10H1-C | - | 8 | GJ211936.1 | 48 | homogeneous | modern | pseudo | 116 | RM |
| 12 | 10 | S1C10H2 | - | 18 | GJ211930.1 | 249 | homogeneous | modern | pseudo | 340 | RM |
| 13 | 12 | S1C12H1L | D12Z3 | 8 | GJ211954.1 | 2350 | homogeneous | modern | live | 35,979 | [10] |
| 14 | 12 | S1C12H2 | - | 18 | GJ211949.1 | 47 | homogeneous | modern | pseudo | 221 | RM |
| 15 | 12 | S1C12H3d | - | 8<sup>6</sup> | AEKP01211346.1 | 23 | divergent | archaic | relic | - | this paper |
| 16 | 10,12 | S1C10/12H1d | - | 2 | ABBA01049496.1 | 93 | divergent | modern | relic | - | this paper |
| 17 | 16 | S1C16H1L | D16Z2 | 10 | GJ212051.1 | 1928 | homogeneous | modern | live | 38,243 | [11] |

<sup>1</sup> A RM is supposed to be a model of a HOR cluster constructed under the assumption that all copies form a single array in one chromosome. For known special cases such as double or triple HOR domains (same live HOR in two or three chromosomes; see section 2.1), the size of RM is adjusted appropriately to represent only one chromosome. For divergent HORs not represented by RMs, only the names of sample contigs are provided. These contigs do not contain all copies of respective HORs and their size does not reflect the copy number.
<sup>2</sup> Genomic size is an estimated length that this HOR occupies in haploid genome. For homogenous HORs represented by RMs it is just the RM length. For S1C1/5/19 RM the size reflects the length of the HOR array on one chromosome under the assumption that the arrays in all three chromosomes are of equal size. For divergent HORs it is calculated from the data in Table 2 (the sum of corrected copy numbers for all monomers of a HOR multiplied to the monomer length).
<sup>3</sup> Pseudo and relic are the two kinds of dead HORs we here propose to discriminate. As a rule, the live and dead HORs can be discriminated by CENP-A binding and the large size of the live arrays (see other columns in this Table). Dead pseudo HORs are homogenous (divergence 1-3%) and dead relic HORs are divergent (9-15%).
<sup>4</sup> The figures shown in this column are the numbers of the 99 bp sequence reads corresponding to a given HOR out of 1 million read sample of CENP-A CHIP-seq dataset (SRR1561921) obtained from a HuRef lymphoblastoid cell line [12]. The whole dataset (about 6 million reads split in 1 million portions) was annotated by HumAS-HMMER used the same way as described in this paper. As the portions did not differ significantly, the data for only one of them are shown in the Table. For S1C1/5/19, the number is adjusted to represent the length of this HOR array on one chromosome under the assumption that the arrays in all three chromosomes are of equal size.
One can see that the live HORs are the major CENP-A binding sites. The data are shown only for homogeneous HORs. The numbers for divergent HORs were slightly higher than for dead homogeneous HORs and much lower than for live homogenous HORs. However, these numbers were not deemed reliable due to admittedly less specific recognition of divergent HORs and more effect the false recognition has on their quantification, both of which could be exacerbated by the short length of the monomer fragments in deep sequencing reads. Thus, the CENP-A binding with these HORs was likely to be somewhat overestimated.
<sup>5</sup> HOR S1C3H2 is also represented in the assembly by GJ211867.1 (length 14 kb) which upon thorough study was disqualified as a valid RM for this HOR (see section 2.2.1 and Supplementary note 1). HOR S1C1/5/19H1L is also represented in the assembly by GJ212205.1 (length 340 bp) and GJ212206.1 (length 340 bp) which were disqualified as valid RMs for this HOR due to their short length (see section 2.2.1 and Supplementary note 1).
<sup>6</sup> In fact, S1C12H3 is not an 8-mer, but a variable size HOR 1-2-3-(4-5)*n*-6-7-8 based on 8 types of monomers (see Supplementary note 1).

### 2.2. SF1 HOR RMs in hg38 assembly. Conceptual overview of analysis performed in this paper

In this section we provide a conceptual overview of analysis performed and tools developed which we report in this paper. In further sections, we present a more detailed description of important stages and a technical description (section 2.13) which lists programs, scripts and settings used.

#### 2.2.1. Non-redundant list of SF1 RMs

For this work we performed a comprehensive phylogenetic and genomic analysis of SF1 as it was represented in the latest hg38 genome assembly. The main new feature of this assembly was that the large centromere gaps were filled in by AS RMs which were approximate representations of AS HOR arrays that formed human centromeres [13]. Initial evaluation and critique of RMs was performed in our recent work [1] which presented the first genome-wide annotation of AS SFs. This paper should be consulted for the brief overview of AS organisation in hg38 assembly, the arrangement of RMs and for the full details of the two main problems uncovered in the organisation and structure of RMs in the assembly. The first concerned redundant sets of RMs in centromeres of acrocentric chromosomes 13, 14, 15, 21 and 22, and the second, a number of erroneously assembled models which featured the reverse order of monomers in a HOR. However, these problems did not involve SF1, so they would not be further mentioned here. The non-redundant list of RMs reported in [1] had 16 SF1 models of which 2 were just 340 bp long and were upon later examination disqualified as models representing a valid independent HOR (see Supplementary note 1, section 1.2.2). Another two RMs were listed as SF mixes containing both monomeric classes characteristic of SF1 (J1 and J2) and of SF5 (R1 and R2). One of these (GJ211866.1 containing HOR3-2 [1]), a dead HOR family located in pericentromere of chromosome 3, was extensively studied [1] and demonstrated to represent the early generation of SF1 AS which we there called “early” SF1 and in this work preferred to term “archaic” SF1, as opposed to “modern” SF1 in most live centromeres. We also defined the J1 and J2 “haplotypes” which listed the single nucleotide differences between the consensus J1 monomer class and its ancestral type A, or the differences between J2 and its ancestral type B. It was shown that archaic monomers had only incomplete J1 and J2 haplotypes while modern monomers had complete haplotypes. Therefore, archaic SF1 monomers were evolutionarily half-way between SF5 and modern SF1. Further analysis (Supplementary note 1, section 1.1.6) showed that the second archaic SF1 RM (GJ211867.1; length 14 kb) listed in [1] lacked a regular HOR and contained pieces of monomers characteristic of the first archaic model. So, it probably represented the same HOR as in GJ211866.1 or rather its shorter variant, but arranged in a scrambled way much like in reverse-assembled models. So, it also had to be disqualified as a model representing a valid HOR. Thus, the SF1/SF5 mixed HORs should be counted as archaic SF1 [1], and a total of 13 valid SF1 models is present in the assembly (see the list in Table 1).

#### 2.2.2. HMMER platform

These models were used as an initial HOR list for annotation. The HORs were extracted and split into individual monomers. The monomers were aligned and after extensive phylogenetic analysis used to create profiles for HMMER [14, 15] as described in full detail in following sections. HMMER package and nhmmer program (for nucleotide hmmer) in particular are the tools for mapping sequence elements in DNA using comparison between the appropriate target sequence region and a number of standard sequence elements installed as profiles in HMMER (termed profile HMMs for “profile hidden Markov models”) [14, 15]. Profile HMMs are statistical models of multiple sequence alignments, or even of single sequences. We also used the consensus class monomers representative of AS SFs [1] as HMM profiles which competed with profiles for individual HOR monomers (see below).

#### 2.2.3. New names

About a half of SF1 HORs identified in the assembly have not been properly described before and had no names, and the ones which had were named in a not very convenient way, as these names only stated that the locus belonged to a repeated sequence in a certain chromosome (e.g. D3Z1 for repeat locus #1 in chromosome 3). These names made no difference between centromeric and non-centromeric repeats and between AS and other centromeric sequences. Also, identical or near-identical sequences situated in different locations had completely different names (e.g. D13Z1 and D21Z1 would represent the same AS HOR located in 2 chromosomes). Therefore, we proposed a new naming system designed only for AS HORs where each HOR received a name which showed its SF, chromosomal location and index number (e.g. S1C13/21H1 for SF1, chromosomes 13 and 21, HOR#1). We hope it would help to keep the information on a large number of AS HORs and individual monomers in a more orderly fashion. The complete rules for the new naming system are listed in Section 2.3 below.

#### 2.2.4. UCSC Browser custom track (the HOR track) and initial evaluation for coverage completeness

The results of initial assembly annotation with this first version of HumAS-HMMER were presented as a UCSC Browser custom track, which was termed the HOR-track. This track was examined in parallel with the SF-track made using the SF module of PERCON program [1, 16], which showed what SF each AS monomer belonged to. SF1 monomers and their arrays were identified in the two-track view and examined for completeness of coverage by monomers that belonged to specific SF1 HORs. The HOR-track was made in such a way that individual AS monomers highly identical to specific HORs were displayed as monomers of these HORs (covered). Individual SF1 HORs were color-coded and the name of each monomer of a HOR could be seen in the full-view UCSC Browser mode (e.g. S1C12H1L.1 for the first monomer of the HOR, S1C12H1L.2 for the second and so on). The monomers which were not similar to any of the HORs would identify with J1 and J2 consensus class monomers the same which PERCON used for SF-annotation [1]. The coverage by class-specific monomers was made invisible in the track, so that respective monomers in the assembly would appear as not covered. The monomers of other AS SFs also identified with their respective class-specific monomers and appeared as not covered in the SF1 HOR-track. Thus, the sequences not covered in the SF1 HOR-track could be: (1) not AS, then they would also not be covered in the SF-track; (2) AS, but not SF1, then they would be covered by a different SF in the SF-track; and (3) SF1, but belonged to an unknown HOR, then they would be covered by SF1 in the SF-track. As a rule the RMs appeared well covered in the HOR-track, but outside RMs we identified several significant arrays which were not covered or sparsely covered in the HOR-track. They were subjected to further analysis in the same way as it was done for RMs.

#### 2.2.5. Evaluation for coverage correctness, color separation and numerical order in the HOR-track

The initial HOR-track was also examined for the correctness of coverage. SF1 RMs were expected to be formed by monomers of the same HOR and hence to have the same color in the track. Indeed, there were only scattered single-monomer imperfections, where presumably erroneous coverage with a different HOR resulted in color mixing. Such “color contamination” cases were largely rectified by introduction of additional monomer profiles in a special cyclic procedure of “color separation” (see section 2.4). Another kind of color mixing which was noted outside RMs was a “mosaic color” where monomers of several different HORs succeeded each other in a chaotic order over a certain region. This was likely to be erroneous coverage especially in the cases where these several HORs were not closely related. Therefore, large mosaic regions were extracted and analysed in the same way as RMs. All erroneously covered and not covered regions appeared to contain not classic homogenous and highly regular HORs, but regions of so-called HOR-like structure previously noted and defined in our recent paper [1].

#### 2.2.6. HOR-like structure regions and relic divergent HORs

The HOR-like structure arrays did contain few distinct classes of monomers similarly to regular HORs, but the classes were much more divergent and their order in an array was much less regular. As a result, the HORs in such regions could rarely be identified by dot-matrix analysis and required full-fledged phylogenetic analysis using trees of aligned monomers. These regions were perceived as old and divergent HOR arrays which have not been homogenized for a long time and have accumulated a lot of mutations (hence divergence) and deletions (hence irregularity). They could be called dead relic HORs as opposed to dead pseudo HORs. Relic HORs are divergent and degraded remains of actual centromeres of our ancestors and pseudo HORs are the results of occasional re-amplifications of pieces of dead arrays. Such re-amplified HORs are homogenous, but often harbor divergent internal repeats within a HOR. Altogether, four divergent relic SF1 HORs were discovered. Some resided only in one chromosome and some in multiple pericentromeric locations. The aligned sets of individual representative monomers for each monomer in each divergent HOR were collected and used for HMM profiles. Such profiles we called MSA (Multiple Sequence Alignment) profiles as opposed to regular single-monomer profiles used for homogeneous HORs. These new profiles were added to the initial set. Then, a new HOR-track was obtained and examined again for the completeness and correctness of coverage. These cycles were repeated until satisfactory coverage of all SF1 sequences was achieved in the final version of the HOR-track. The above described characteristics of all SF1 HORs (live vs. relic vs. pseudo and homogenous vs. divergent) are summarized in Table 1.

#### 2.2.7. No new homogenous HORs discovered

All homogenous SF1 HORs identified in hg38 assembly were previously known to researchers. Some were described in detail, some only briefly mentioned. For some of them, no information was published in papers that we know of, but they were used in various computer programs and HOR lists circulating in the community. In some cases, they appeared for the first time as AS RMs in hg38 genome assembly. No homogenous HORs not represented as RMs were discovered in SF1. In a lot of cases, it was hard to tell what HOR was mentioned in a paper as the only information given was the length of HOR and location. One has to keep in mind that the length of the same HOR can often be stated differently as in each case another structural variant of a HOR may be judged to be the major constituent of an array. HORs would be much easier to identify if their SF was always stated along with the length and location. Therefore, we included SF in the name in our proposed new naming system. We listed the principle references to SF1 HORs that we knew of in Table 1 along with the old names of HORs, if any. This list of references, however, is bound to be incomplete for above given reasons. The 4 divergent SF1 HORs were first described in this paper.

#### 2.2.8. Phylogenetic data on the HOR monomers

Phylogenetic analysis of the SF1 monomers mainly gave two kinds of data that were collected and examined. The first was the information on sequence relationships of monomers within and between HORs (noted in Fig. S1, S2 and Supplementary note 1). The second was the distribution of HORs into 2 broad groups, archaic and modern SF1, which was clearly seen on the trees of all J1 and J2 monomers used as HMM profiles (Fig. S2b) and was confirmed by subsequent haplotype analysis performed as described earlier [1] (Table S1). These results are presented in more detail in sections 2.6, 2.7 and 2.10 below.

#### 2.2.9. Statistics of HOR structural variants in RMs

The annotation obtained with the final version of the HOR-track allowed collection of genomic statistics for SF1 HORs. This genomic analysis was done solely for RMs, where they were available, as they all came from one person (HuRef) and were all in direct strand which simplified the analysis. Besides, RMs are supposed to represent the genome-wide copy number of respective HORs and the HOR copies which appear in the assembly outside RMs would be counted twice. In cases of divergent HORs, which had no RMs, clones or assembly regions which had high concentration of these repeats were analysed instead of RMs in the same way. In these cases, the analysis was not genome-wide and reflected only the HOR pattern and copy number in a specific locality. Notes to that effect were made in respective pages of Fig. S1.

Our analysis yielded HOR maps of RMs and statistics of different structural variants of HORs in RMs. Note that different HORs appeared to differ greatly in diversity and number of structural variants. Some were non-polymorphic (had no or almost no variants) and some were very polymorphic (had a lot of variants interspersed in an array). As it is known that individual humans (and individual chromosomes) differ in concentration and distribution of certain HOR variants, in the future, HumAS-HMMER may be instrumental in personalized analysis of HOR variants [17] which has been shown to be crucial for detection of functional centromeric epialleles and aberrant poorly functional centromeres [4, 5].

Below we describe in more detail all stages of this work.

### 2.3. The rules of the new HOR naming system (examples may be fictional)

The rules provided below cover not only SF1, but also the structural features in other SFs, which do not occur in SF1. We chose to provide the complete set of rules to avoid a need to publish slightly different versions with HOR-tracks for SFs 2-5 later on.

1. The name of the HOR shows SF (S), the number of a chromosome where the HOR is located (C) and the arbitrary number of the HOR in this location (H) (e.g. S1C1H1 means SF1, chromosome 1 and HOR#1 in chromosome 1). These main parts of the name may be used with optional additional indices to highlight useful information about the HOR. Distinct sequence variants of a HOR (as opposed to structural variants) which are segregated in separate arrays have the same name, but with a different letter index given with a hyphen (e.g. S5C1H2-A and S5C1H2-B). Structural variants (almost identical in sequence, but distinct in structure) of the same HOR (e.g. deleted derivatives) are not given separate names, but are treated as variants of the same name. These variants are given arabic numbers (#1, #2, etc.) which are not part of the name and their structure is shown in Fig. S1. Note that sequence variants may also be structurally different (see below).
2. SF is shown according to PERCON [1] and the SF-track, but the SF1/5 and SF2/5 mixes, which are now known to be archaic SF1 and SF2 are shown as the respective new family.
3. Chromosome location may show more than one chromosome with a slash (e.g. C13/21). That means that the same HOR is found in two different locations.
4. If there are multiple locations one can use the CM symbol (Chromosome Multiple).
5. If convenient, additional specification of chromosomal location is possible (e.g. C1p for the short arm and C1q for the long arm side of the centromere).
6. The arbitrary number of a HOR is given irrespective of the SF. Thus, there will be S1C1H1, S3C1H2 and S5C1H3, etc.
7. The number of the single live HOR in a given chromosome is always H1 and additionally marked with a letter “L” to make it easily identifiable (e.g. S1C1H1L). However, H1 does not always has to be a live centromere (see below).
8. Complex locations like C13/21 or CM have their separate numbering. If there are S1C1/5/19H1L and S5C5/19H1, the L index is crucial to identify a live centromere.
9. If there are few sequence variants of the live HOR, the L index is used instead (not together) of the -A index. Thus, in chromosome 17, there are S3C17H1L, S3C17H1-B and S3C17H1-C. Such reduction of the second letter index occurs only for the live centromere. This is done to save space and make names a bit less complicated.
10. An optional additional index “d” is used to show the divergent HORs, as opposed to homogeneous (e.g. S1C3H4d). Both indices L and d never occur together, as all the live centromeres are made by homogeneous HORs.
11. Different generations of SF1 and SF2 (archaic and modern) may be reflected in the name if needed by using letter indices (“a” and “m”, respectively) with the SF part of the name (e.g. S1aC3H2 for archaic SF1 HOR in chromosome 3). As modern HORs are prevalent, they may go without the “m” index, then only the rare archaic HORs will be marked.
12. Individual monomers of a HOR are shown as H1.1, H1.2, H1.3, etc. according to the numbering of a basic HOR shown in Fig. S1 as variant #1. In different structural variants (#2, #3, etc.), the straight order will always be modified and will show the difference with the basic HOR. In different sequence variants (different letters of the same name), the numbering of monomers is the same as in basic HOR. If the sequence variant is structurally unchanged, the straight numbering is preserved. If in such a variant there is also a structural difference (e.g. deletion or duplication), the order may deviate from the straight (e.g. the basic HOR would have the order 1-2-3-4-5, the deleted variant would be 1-2-4-5, and duplicated variant would be 1-2-3-4-3-4-5). The basic HOR (variant #1 in Fig. S1) would always have the straight order.
13. Some deletions in derived HORs may happen out of register. If one monomer is deleted which includes one half of monomer 2 and one half of monomer 3, a hybrid monomer 2/3 is formed. It has the first half of monomer 2 and the second of monomer 3. A pentamer with such a deletion will appear as a tetramer 1-2/3-4-5. Sometimes the deletion breakpoints would not be in the middle of the monomers. For instance, a hybrid monomer may have one third of monomer 2 and two thirds of monomer 3.
14. If the two monomers within a HOR are highly identical and are not distinguished by HMMER they are given a combined number and appear twice in the HOR (e.g. 1-2&6-3-4-5-2&6-7-8).
15. If an alien monomer which does not originate from the basic HOR, appears instead of a monomer in the derived HOR (or only in some of its structural variants), such monomer is denoted by a small Latin letter (e.g. 1-2-3-4-5 in the basic HOR vs. 1-2/x-x/3-4-5 in the derived variant). Such monomer may also appear as an insertion (e.g. 1-2/x-x/2-3-4-5).
16. If the HOR sequence variants are structurally not simple derivatives of the basic HOR, but are related in a complex way (e.g. some monomers are the same, but some are different, or the basic HOR is divergent and the variant is a homogeneous new amplification of its segment), they were given entirely different names, but their relationship was noted and recorded in Supplementary Note 1 and Fig. S2.

### 2.4. Formation of the HOR list and profile sets for AS annotation

In this work, we employed a “master HOR” approach to annotation of homogenous AS HORs. We picked a single HOR copy from each RM in our non-redundant list of 13 SF1 RMs described in section 2.1 (see also Table 1), split it into monomers and installed each of them as a separate HMM profile. The process of selection of such a “master HOR” copy is described in section 2.5. Such sets of HOR-specific profiles were used along with a set of SF-specific monomer class profiles described previously [1]. These were the profiles for J1, J2, D1, D2, W1-5, R1, R2 and M1+ classes of monomers used as they were published in [1]. Then, a track for the chromosome from which the RM came was made in such a way that the coverage by the HOR monomers was visible and the coverage by SF-specific profiles was invisible and appeared as not covered. Such track was examined for completeness of coverage in the respective RM which contained the HOR and in possible copies of the HOR in genomic clones outside the RM. Examination was performed in parallel with SF-track [1] which visualized both HOR-covered and not covered AS. As expected, the HOR monomers tended to cover the HOR arrays, but the other sequences of the same SF appeared as not covered (in fact covered by invisible SF-specific class monomers). If the RM coverage was not complete or nearly complete, the monomers within the RM which were not covered or covered with a different type of HOR (erroneous coverage which appeared in the track as mixing of colors) were collected, marked according to their position in the HOR and introduced as additional profile monomers. The names of such additional monomers were identical and the same as the name of the original master HOR monomer. For instance, the uncovered or erroneously covered monomer that was located between monomers 4 and 6 in the HOR would be tentatively marked as monomer 5 and its identity would be verified by phylogenetic analysis. This process, which we termed “color separation” procedure was reiterated to resolve imperfections in the tracks until complete or near-complete correct coverage was achieved.

For many HORs (non polymorphic ones) a single master copy of a HOR was sufficient. For others (polymorphic) we had to add many monomers to get a good coverage. This manual procedure amounts to rough characterization of a degree of sequence polymorphism of a given HOR and yields a list (albeit perhaps redundant and incomplete at the same time) of different variants of each monomer in a HOR. These lists can in the future be refined and expanded to cover all sequence variants of a certain HOR at the desired level of detail. If the profiles for different variant monomers are marked by different names, the program would annotate different sequence variants (monomer or HOR haplotypes) in the array. So, it can be developed into a powerful tool to study sequence polymorphism in AS HORs. As of now, all polymorphic variants of the same monomer were marked the same, and no detection of sequence polymorphism within the same HOR was enabled. However, we were able to observe structural variants in which numerical order of monomers differed from the master HOR (deletions, substitutions and duplications of monomers and more complex variants such as hybrid monomers). Also, sequence variants of a HOR which were segregated into different arrays were noted and labelled with different hyphenated letters of the same name (e.g. S1C10H1-B and S1C10H1-C).

Note that a special care was taken to choose the master HOR in such a way that it represented the most regular and simple and in most cases the most abundant variant (variant #1) which may also be called a “basic” HOR. Such HOR features a basic order of monomers (straight) and all or most other HOR variants are presumably derived from it via some easily discernible event like deletion or duplication. In practice, however, it is not always the way and the origin of some variants could be rather more complex. To quantify common variant HORs in RMs (which all come from the same HuRef individual and all have AS in direct strand) we constructed simple RM maps in which listed all monomers, and where a single HOR copy would occupy one line. A new line would begin every time monomer number 1 occurred in an array. Such maps for all RMs were attached to Fig. S1 (as indicated by the links), where the statistics of most abundant variants was also shown. Full statistics can be viewed via the links provided. In some cases, where a HOR (or a sequence variant of a HOR) was not represented as RM (see below) we made the maps of genomic clones or contigs where the HOR was present. Some of such clones had AS in reverse orientation, therefore, in the maps, the order of monomers should be read in reverse order (from right to left), and monomer number 1 of a HOR appeared at the end of a HOR instead of a start. Such maps were marked with a special note to that effect.

All RMs which were SF1 according to the SF-track were processed in the above-described way one by one, each adding sequentially to the number of profile monomers used in the working version of HumAS-HMMER. Initially, for each HOR, only the chromosome(s) in which the HOR was known to reside was studied. When good coverage for all SF1 HORs was achieved, the track for the whole assembly was made and re-analysed and color separation procedure was applied as needed.

As SF1 is one of the new SFs, it is not supposed to have a non-HOR component [1, 3], and complete coverage was expected for all regions in the assembly that were SF1 according to the SF-track. Therefore, if some such region was not covered or was covered erroneously in the HOR-track, it was likely formed by some new HOR that was not represented by a RM. If the HOR could be revealed by the dot-matrix, it was picked up in a standard way and added to the list. If not, such regions were extracted as a whole and their monomers aligned and analysed using phylogenetic trees. If a few distinct branches (divergent classes of monomers) were observed, the locus was classed as a HOR-like structure [1]. Then, the monomers from each branch were collected and used for HMM profiles as MSA. In the track, the consensus order of monomers in the new HOR was established, the maps of respective regions constructed and the variants catalogued according to the established procedure. After the new HORs were added to the list, the combined track for SF1 was made again to ensure the complete coverage. It has to be noted that color separation in the track for divergent HORs (the ones that had MSA profiles) was often worse than for homogenous HORs (the ones with single-monomer profiles). For instance, the S1CMH1d arrays are often contaminated by S1C3H3d monomers. Some of these contaminating monomers may be real S1C3H3d insertions, but some are likely due to erroneous coverage. As the profiles for divergent HORs were MSA, we could not use our standard color separation approach to address impurities as addition of single monomers to the profile would not change it significantly. Thus, some more sophisticated methods were called for. At this point, we just accepted that the resolution for divergent HORs was not as good as for the homogenous ones.

We also took special care to make sure that for all the monomers marked as certain monomers of a certain HOR and used for HMM profiles their position on a monomer tree was consistent with their assignment. For this purpose, for each HOR we considered: (1) a tree which had only the monomers of a master HOR (only one copy of each monomer of a HOR); (2) a tree having all the monomers used to identify this HOR (some monomers may have multiple copies); and (3) a tree having all the monomers used for HMMER of the same SF class. The first and the second trees for each HOR are shown in Fig. S1 and the third tree is shown in Fig. S2b. The details of this procedure are described below.

### 2.5. Typical processing of a RM or a genomic region

The typical analysis of a RM was performed as described below. If a HOR was not represented by a RM, a genomic clone or a traditional contig which contained the HOR was picked up and analyzed in the same manner.

The master HOR was selected randomly as a typical copy of a HOR as follows. A dot-matrix of an RM was examined and a region with a clear HOR structure was selected. From this region a sample of sequence containing 3-5 copies of a HOR was extracted and used for alignment. The monomers were extracted by PERCON and their SF monomer classes were established (shown in Fig. S1). Position 1 in the monomer by tradition was arbitrarily assigned to the first nucleotide of the BamHI site in chromosome X-specific HOR DXZ1 [18]. The aligned copies of a HOR were compared and one copy that lacked irregular features (e.g. deletions or insertions which were present in some copies and absent in the others) was selected as master HOR. Its monomers were used as profile monomers in HMMER, HOR track was prepared and analysed for completeness of coverage, and additional monomers were added to achieve complete or near-complete coverage by respective HOR, as described above.

When a good coverage was achieved, the RM was examined in a combined track where all HORs of the same SF were installed and the color separation within the SF was examined. Note that in our work the HORs were added to annotation sequentially, so the first ones added had really no partners to mix with and the last ones had a full set of partners. Thus, at the end, the whole issue of color separation had to be re-analysed anew. Similarly, after additional monomers were introduced at any point, the color separation in one locus could be improved, but in the other could become worse. So, the whole track should have been monitored constantly. The current level of separation is a compromise and may be improved in the future. For sequence variants of the same HOR segregated in separate arrays (different hyphenated letters of the same name) even a significant degree of mixing was considered tolerable given that one of the colors was clearly dominant. An equal or near equal color mix would suggest that the two HORs are the same and should be treated as such. Thus, the degree of color mixing was even considered a useful measure showing just how close the sequence variants of the same HOR were. If no significant separation could be obtained with a limited effort, such HORs would be considered one and the same (if structurally identical) or structural variants of the same HOR (if different). Theoretically it is possible that the process used to construct RMs could artificially segregate into different RMs the variants which are actually mixed within the same array. So, groups of closely related HORs like S1C10H1L, S1C10H1-B and S1C10H1-C should be further studied and their segregation in traditional genomic contigs has to be demonstrated. We believe that such studies would be well assisted by the use of the HOR module of HumAS-HMMER we present here.

The monomers used for the profiles were analysed to make sure their name assignment did not contradict their position on a tree. The trees for monomers of a master HOR (first tree), for all monomers of a given HOR (second tree) and for all monomers of SF1 (third tree) were constructed and examined sequentially for consistency. The first and second trees for each HOR are shown in Fig. S1. Relationships of monomers within a HOR were noted on the first tree (marked by color in Fig. S1 and noted in Supplementary note 1). On the second tree, we made sure that all somewhat different copies of the same monomer branched together and hybrid monomers which occupied positions different from positions of a master HOR monomers were noted. On the third tree (Fig. S2) we made sure that no monomers labelled by different names were in fact identical or closely related. Relationships between HORs were also noted on the third tree and summarized in Fig. S2 and Supplementary note 1. Also, correctness of the monomer classes established by PERCON was additionally confirmed as monomers of the same class should branch together. Moreover, the hybrids between classes were identified on the second and third trees due to their unusual position (between classes).

Finally, the monomer map of each RM was constructed and the statistics of structural HOR variants gathered. Also the statistics for SF monomeric classes (PERCON classes) for each monomer of a HOR was collected to make sure that PERCON assignments in Fig. S1 showed the typical pattern. This stage yielded a catalog of structural variants present in a RM for each HOR. The links to the maps are provided in Fig. S1. Programs and scripts used are listed in technical description (section 2.13).

Note that the identity of a monomer may be established not only by its coverage but also in a tentative manner by its position in an otherwise well-covered HOR. For instance, if the monomer positioned between the 4th and the 6th monomer of a HOR is often not covered or covered erroneously (by a monomer of another HOR), one would suggest that there is a polymorphic variant of the 5th monomer, which needs to be added to the set. Indeed, the erroneous coverage was often consistent, so that this 5th monomer was mostly covered by, for instance, the 2nd monomer of such-and-such another HOR. However, there was always a theoretical possibility that such a case was not really an erroneous coverage, but a genuine one and a monomer of the second HOR really occurred in an alien genomic context. This could be checked by looking at the scores in a track (the scores for genuine coverage of an alien monomer would be higher than for erroneous), or on the trees where the monomer in question would branch either with the fifth monomer(s) of the first HOR (erroneous) or with the 2nd monomer(s) of the second (genuine). In our practice, the color mixing in the tracks was mostly caused by erroneous coverage. In this work, we made only limited effort to improve color separation by adding profile monomers, focusing mostly on mixing between different HORs. Mixing of sequence variants of the same HOR was considered tolerable, and moreover a useful measure of identity between the variants. Similarly we adopted reduced requirements to color separation in divergent HORs. Note that our system of naming and numbering was especially designed to allow reading the HOR structure in the track despite mixing between sequence variants (different hyphenated letters of the same name).

The total number of copies of each monomer in each divergent SF1 HOR was calculated from the HOR-track using the Table Browser. It was corrected by subtracting the number of falsely recognized monomers calculated as the sum of the monomers of a given HOR which were SF2 (total of 8), SF3 (total of 1) or SF4+ (total of 180) according to PERCON (the SF track). It was not possible to correct for the false recognition of SF5 monomers as all SF1 divergent HORs except for S1C10/12H1 were archaic and PERCON recognized archaic SF1 as SF1/SF5 mix [1]. S1C10/12H1 was not significantly involved in false SF5 coverage (the total of false SF5 hits was 1).

### 2.6. Processing of divergent and homogenous HORs

Divergent HORs were discovered as not covered or erroneously covered SF1 regions where classical regular HORs were not observed using dot-matrix analysis. When such regions were extracted and the monomers aligned and analyzed by construction of phylogenetic trees, we observed few kinds of monomers as in regular HORs. However, monomers of the same kind were not nearly identical (1-2% divergence), as in classical homogeneous HORs, but differed on average by 9-15%. When different kinds of monomers were marked and maps of the arrays made, a semblance of numerical order typical of the HOR arrays was observed albeit much less regular. Thus, it was obvious that these were the old diverged HORs partially ruined by deletions and perhaps some other recombination events frequent in tandem arrays.

To compare these novel divergent HORs with classical homogeneous ones we calculated the divergence for each monomer of the 4 divergent HORs using the aligned monomers obtained for our analysis (monomers are shown in Alignment file 1) and for the monomers of the 4 selected homogenous HORs. For homogeneous HORs, the sample regions containing about 300 HOR copies were randomly picked from respective RMs. The results are shown in Fig. 1 and Table S2. One can see that less polymorphic homogenous HORs S1C7H1L, S1C12H1L and S1C16H1L had the classical degree of divergence of 1-2%, and more polymorphic homogenous HOR S1C10H1L had slightly elevated divergence of about 3% (see Discussion) while divergent HORs quite deserved their name being 9-15% different.

**Figure 1.**
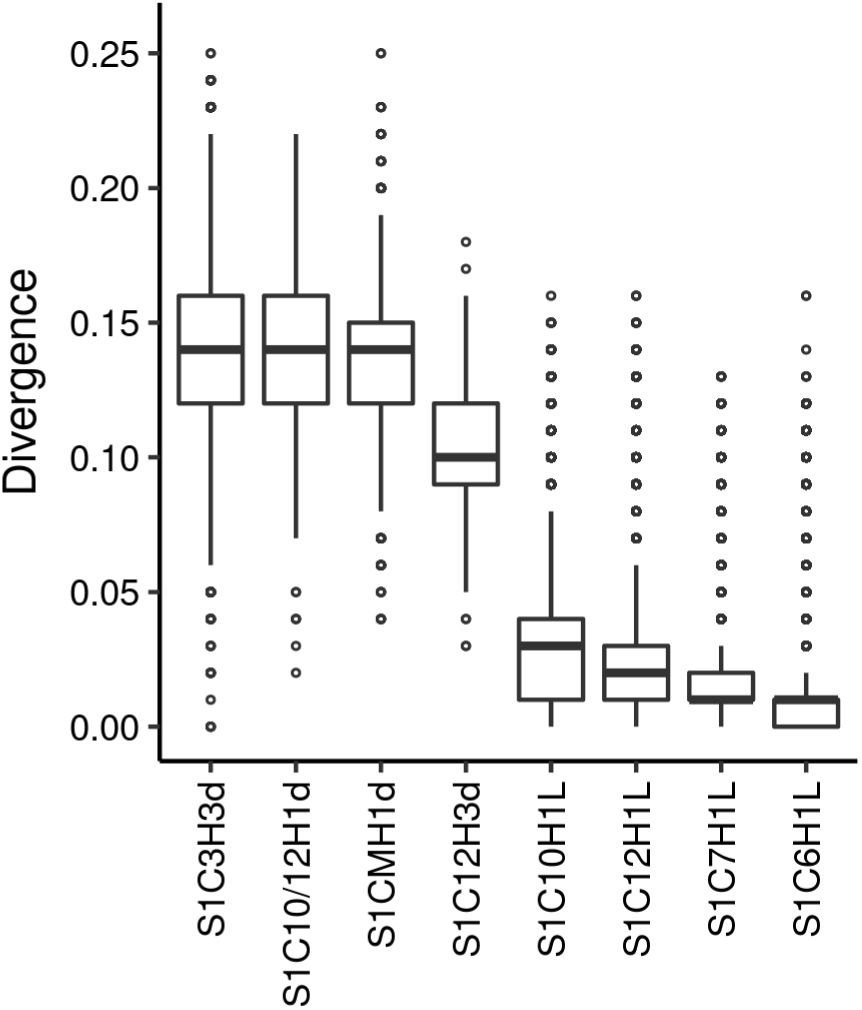
Divergence of selected SF1 HORs. This boxplot displays the difference in homogeneity between divergent and homogenous HORs. It visualises five summary statistics (the median, two hinges and two whiskers), and all “outlying” points individually, as described at (http://ggplot2.tidyverse.org/reference/geom_boxplot.html). For divergent HORs (marked with letter “d” in the name), divergence (%) was calculated using the sets of aligned monomers shown in Alignment file 1. The number of monomers and the divergence in each set are shown in Table S2. As the figures for different monomers of the same HOR were pretty consistent, the divergence values were pooled for each HOR and used for this generalized boxplot. Four different homogenous live HORs (marked with letter “L” in the name) were used for comparison. One of them was highly polymorphic (S1C10H1L) and three were non-polymorphic (S1C7H1L, S1C12H1L and S1C6H1L). For each, a region in respective RM containing 200-300 copies of each monomer of the basic HOR was picked up and individual monomers with a length 150 bp or longer were extracted by the Table Browser using their HumAS-HMMER assignments and aligned. Only the monomers of a basic HOR were used for comparisons. The hybrids were not included, although they were present in the sample regions, as we suspected there could be few different kinds of each hybrid which resulted from independent deletions or other events. In such case, few kinds of each hybrid could exist and hybrids would not be expected to be homogenous. Therefore the sample region for the polymorphic HOR had to be much larger than for the non-polymorphic, as a large number of hybrids occurred in the former. One can see that there is a large gap in divergence between homogenous and divergent HORs. See the note to Table S2 for more details.

The copy number of homogeneous HORs is probably best represented by the lengths of respective RMs [4] (shown in Table 1) as RMs are supposed to represent the genome-wide copy number of respective HORs. Thus, for regular chromosome-specific HORs which are all gathered in one array this should be a good estimate. For the HORs present on more than one chromosome like S1C1/5/19H1L, the RM estimate may not be so good, as it assumes that each chromosome has the same number of HORs which may not be true. The copies of a HOR that may be present outside RM theoretically should not be included in the estimate as they will be counted twice, but in practice this number is usually too small and therefore insignificant. Few HOR monomer hits that may appear over an alien SF as a result of possible false recognition would be even less significant (see section 2.8 below). In any case, all such estimates are very approximate as individuals differ widely (up to 10 times difference) in the size of AS arrays [13, 19]. Thus, non-personal estimates are only good enough to indicate the order of magnitude for the copy number. Note that RMs are all derived from the same individual (HuRef) and the precise RM-based estimates apply to this individual only. The copy number of divergent HORs (Table 2) which are not represented by RMs can best be estimated from the HOR-track, but in the case of these extremely low copy number HORs false recognition may potentially be significant and should be controlled for. In Table 2, the total number of copies for each monomer of each divergent HOR is shown and corrected for the number of the same monomers which were found over D1, D2, W1-W5 and M1+ monomers (false coverage). Of these, only the coverage over M1+ (SF4+) monomers was significant. Note that we were not able to correct for the false coverage over SF5, because a lot of monomers in archaic divergent HORs (all but S1C10/12H1) were classed by PERCON as R1 or R2 monomers. However, the manual inspection of the tracks indicated that the amount of false coverage over the true SF5 arrays was comparable to that over SF4, and could not greatly affect our estimates.

It was clear after initial experiments that annotation of divergent HOR regions could not be achieved with the “master HOR” approach we used for homogeneous HORs, where a single monomer or just a small number of profile monomers were sufficient to ensure a good coverage. With this approach we observed only partial coverage in divergent arrays where many monomers appeared to identify with SF-specific monomer class profiles and were not visible in the track. Instead we used a “master region” approach where a large number of monomers of the same kind from a single region or from few regions were aligned and this multiple sequence alignment (MSA) was used as HMM profile, which in this case we called a MSA profile. Experiments revealed that such profiles provided not only for satisfactory coverage of the region from which the alignment was made, but also of similar regions in different loci on the same or on the other chromosomes. Such regions were covered with the same or similar numerical order as in the “master region” and thus were considered the arrays of the same HOR. Obviously, in case of divergent HORs, discrimination of slight sequence variants of the same HOR using color separation was not likely, as it was largely masked by high sequence divergence and only structural variants could be observed. Also the quality of coverage was on average poorer than in homogeneous HORs.

**Table 2.** Copy number estimate for divergent SF1 HORs.

| **HOR name** | **Total monomers** | **Corrected total** |
| --- | --- | --- |
| S1C10/12H1d.1 | 358 | 259 |
| S1C10/12H1d.2 | 288 | 288 |
| S1C3H3d.1 | 251 | 248 |
| S1C3H3d.2 | 237 | 230 |
| S1C3H3d.3 | 318 | 310 |
| S1C3H3d.4 | 247 | 243 |
| S1C3H3d.5 | 259 | 239 |
| S1C12H3d.1 | 8 | 8 |
| S1C12H3d.2 | 9 | 9 |
| S1C12H3d.3 | 11 | 10 |
| S1C12H3d.4 | 50 | 36 |
| S1C12H3d.5 | 38 | 37 |
| S1C12H3d.6 | 11 | 11 |
| S1C12H3d.7 | 11 | 11 |
| S1C12H3d.8 | 14 | 14 |
| S1CMH1d.1 | 395 | 382 |
| S1CMH1d.2 | 512 | 505 |
| S1CMH1d.3 | 366 | 356 |
| S1CMH1d.4 | 228 | 226 |
| **Total** | **3611** | **3422** |

### 2.7. Processing of hybrids

Only a limited effort to identify hybrid monomers was made. Specifically, (1) if the hybrid nature was pretty obvious upon inspection of the aligned monomers of the same HOR, or (2) if a monomer had an unusual position in a tree, or (3) if the hybrid nature was revealed by monomer context, or (4) if the trees made of the right halves of the monomers and the left halves were different, was a monomer analyzed to establish its hybrid nature. The hybrid status was assigned only if the nature of the monomer was pretty obvious. For example, a common HOR variant 1-2-?-7-8 (where “?” is a monomer unusually positioned in a tree) suggests that the “?” may be a 3/6 hybrid formed by an out-of-register deletion of monomers 3-6 (3 and 6 deleted partially, 4 and 5 completely). On the other hand, many hybrids between more closely related monomers or the ones which have only a small piece of one monomer and a large piece of the other, may have gone unnoticed because in alignment and in the trees they look pretty much as monomers of a basic HOR. Some data on the validity and copy number of specific hybrids is provided in Supplementary note 1, section 1.2) As an example, we have particularly studied the genomics of “cen1-like” AS dimer published by Henikoff et al. [12, 20], which appeared to be S1C1/5/19H1L (6/4-5) dimer according to our monomer numbering (see section 1.2.2 in Supplementary note 1 for details).

### 2.8. The background of false SF1 coverage in non-SF1 arrays

In a number of regions in the assembly, we did observe a significant background coverage of SF1 HOR monomers over the arrays which were not SF1 by the SF-track. This background false coverage appeared as constellations of separated single monomer hits which we termed “clouds” as opposed to continuous true coverage over the SF1 arrays. Such clouds almost never appeared over the SF2 and SF3 arrays, but sometimes were observed over SF5 and SF4+. At high magnification in the track it was readily distinguishable from the true coverage, but at bird’s eye view such clouds would look like isles of SF1 sequence. For this reason we recommend to always view the HOR-track in parallel with SF-track which allows easy identification of such false coverage at any magnification. The number of false SF1 HOR hits over SF4+ arrays could be estimated as the number of monomers in the HOR-track which corresponded to M1+ monomers in the SF-track, these made up 0.2% of total SF1 HOR hits. In SF5 arrays, this method was not applicable as archaic SF1 HOR arrays appeared in the SF-track as SF1/SF5 mix. However, for the modern SF1 arrays it could be used, and we estimated that less than 0.1% of total modern SF1 HOR hits were involved in false coverage of SF5 arrays. Thus, the overall false coverage was insignificant, but if one looked only at divergent HORs which were much less numerous than homogenous ones, the background of false hits over SF4+ reached 5%. Manual examination of the tracks suggested that the extent of false coverage over SF5 which could not be correctly estimated in a genome-wide manner was of comparable magnitude. So, the copy numbers for divergent archaic HORs determined from the tracks and shown in Table 2 may be overestimated by about 5%, as they were corrected for false recognition over SF4+, but not over SF5. The hits in the clouds over M1+ usually corresponded only to a part of a monomer and had lower scores than the hits in monomers of homogenous HORs. However, among divergent HOR monomers, there were true hits with comparable HMMER scores, so we did not want to eliminate the clouds by increasing the filtering threshold (score to length ratio threshold set at 0.7; see section 2.13.2 below), as that could decrease coverage in divergent HOR arrays. Our preliminary experiments demonstrated that the proper way to tackle the clouds would be to improve the class profile for M1+ monomers. If separate profiles for all constituent colored layers within M1+ arrays [1, 3, 12] were introduced, the false SF1 background coverage over SF4+ arrays almost disappeared. This set of additional profiles will be described in a separate report and the next generations of the HOR-track will feature much less clouds over SF4+. We suppose that this approach will also be efficient for the false hits over SF5 where R1 and R2 monomeric classes should each be represented by a number of somewhat different profiles. This will be possible upon completion of a more detailed genome-wide study of possible phylogenetic groups of monomers within SF5, which is under way. For now, we chose to tolerate the amount of clouds there is in the track as it does not much interfere with the viewing. However, it may somewhat hamper gathering genome-wide statistics for some HORs, so only the statistics for isolated SF1 genomic segments such as RMs or contigs was used throughout this paper, except for Table 2 (see the legend and section 2.6).

### 2.9. The consensus succession of SF1 layers in centromeres

The succession of different generations of SF1 HORs in a single chromosome can be elucidated by manual examination of the HOR-track despite it being obscured by numerous rearrangements and deletions in pericentromeric regions, and a consensus order can be established. If a live HOR is positioned in the center and the SF5 is on both flanks, the most distal SF1 HOR would be S1CMH1d which could be followed by one or more of the other archaic SF1 HORs, then the modern layer starts with S1C10/12H1d, which may be followed by one or more modern SF1 pseudocentromeres and then by the live SF1 centromere. Note that in some chromosomes archaic SF1 layers may be followed by SF2 live centromere. For instance, in chromosome 8 which has SF2 live centromere, there are chunks of S1CMH1d in both p- and q-arms (chr8:43,936,718-43,966,062 and chr8:45,945,726-45,971,761, respectively) which seem to be segment duplications (SDs) of one another (parts are 96.5% identical). These SDs however are not not reflected in the UCSC SD track (see discussion of AS SDs in hg38 assembly in [1] and [21]). Whether these SDs indicate the former presence of SF1 centromere in chromosome 8, or they just came from another chromosome is not clear. In chromosome 1, the remains of older SF3 centromere [1, 3] surround the live S1C1/5/19H1L array and in this case there are no archaic SF1 layers around the latter, but SF1 live HORs are directly flanked by SF3 pseudocentromeric HORs. Also, in chromosome 16, the remnants of archaic SF2 HOR flank the live SF1 centromere ([1] and our unpublished results).More remarks on the succession of SF1 domains in individual chromosomes are provided in Supplementary note 1.

### 2.10. Haplotype analysis

Haplotype analysis was briefly described in section 2.2 above. It was performed as previously described [1] and the results are shown in Table S1. Slight changes introduced in consensus sequences used were as follows: (1) “N” positions in class consensus monomers were substituted by combinations of letters which showed the actual composition of nucleotides in those positions, according to class matrices published in [1] or to the actual monomer sets for divergent HORs shown in Alignment file 2; (2) J1m and J2m consensus monomers (see below) derived only from modern master HOR monomers (Alignment file 3) were used instead of J1 and J2 consensus monomers used in [1] which were derived from a mixture of modern and archaic monomers; (3) consensus monomers for types A and B derived from consensus monomers of SF-specific classes underwent slight changes due to the changes in constituent class consensus monomers and to the less stringent derivation rule (see below). These slight differences in consensus monomers are summarized in Table S3. Overall, the changes involved only a few split or ambiguous positions some of which became unambiguous in result. All consensus monomers used in this work in their regular form (with “N” positions) are provided in the files with sequence alignments attached to this paper (Alignment file 2 and 3). Consensus monomers J1m and all J2m were derived from the monomers of modern SF1 HORs (monomers of archaic HORs and archaic part of S1C3H1L were left out). To avoid bias towards any specific HOR, consensus sequences were derived from monomer sets which included only the master HOR monomers (for homogenous) and consensus HOR monomers (for divergent). So, all the hybrid monomers were left out. Consensus divergent HORs were derived from the sets of monomers used for HMM profiles. The number of monomers in each set is shown in Table S2. All the HOR and the class consensus monomers used in this paper were derived using the “50% or more” rule. The A and B type consensus versions used for haplotype analysis were derived using a less stringent rule. There, a simple prevalence of a nucleotide among the 5 type A class monomers (J1m, D2, R2, W4 and W5) or among the 6 type B class monomers (J2m, D1, R1, W1, W2 and W3) was sufficient to draw an unambiguous position. Respective class monomers are shown in the upper part of appropriate section in Table S1 right under the type consensus monomer. The Table is divided into two parts according to A and B types. Additionally, in type A positions 15 and 153, where there was no prevalent nucleotide we took into account that most of archaic J1 monomers differed from J1m in these positions. Therefore, the “G” nucleotides which appear in both positions in J1m could not be carried over from ancestral type A monomers, as they were not present in the immediate ancestor of J1m. Thus, they appear to have mutated into “G” recently and independently of the “G” in position 15 of W5 and position 153 of W4 (see the upper part of Table S1, type A). Hence, we discounted these coincidences and designated the other prevalent nucleotides as ancestral in type A (“C” in position 15 and “T” in position 153). Also, while deriving the consensus of type B, we considered the deletions in position 68 of J2m and W3 class consensus monomers to be different deletions, as one of them was a single nucleotide deletion and the other was a part of a three nucleotide deletion in positions 66-68 (see the upper part of Table S1, type B). So, this coincidence was discounted as well.

Results shown in Table S1 demonstrate that monomers of archaic HORs have only partial SF1 haplotypes and the monomers of modern HORs have complete or almost complete haplotypes.

### 2.11. Inter-HOR relationships

Relationships in the group of archaic HORs are pretty straightforward. They all originate from S1CMH1d or a very similar HOR with the same 4 kinds of monomers (Fig. S2a). Only the first 3 of these occur in S1C12H3 which is closest to the progenitor, and all 4 in S1C6H1L which is more distant. Another two archaic HORs more loosely related to the progenitor are S1C3H3d and S1C3H2. They also contain only the first 3 monomers of S1CMH1. S1C3H3 is a divergent pentamer and S1C3H2 is a recent amplification of 2 such divergent pentamers which form a homogenous 10-mer HOR (see HOR 3-2 in [1] for detailed analysis). To the contrary, in the group of modern HORs, which all originate from divergent S1C10/12H1d or similar sequence, no clear relationships are obvious except for S1C1/5/19H1L and S1C16H1L where the first is obviously but imperfectly collinear to the first 6 monomers of the second (Fig. S2a, Group 2). More distant and less clear-cut relationships are observed between the HORs residing in one chromosome which make 3 pairs: S1C1/5/19H1L and S1C5H2; S1C10H1(L, B and C) and S1C10H2; and S1C12H1L and S1C12H2. In all these cases, the H2 HORs tend to be longer and to have more kinds of monomers than the H1 HORs (Fig. S2a). However, it does not seem that any of the H1 HORs could be direct re-amplifications of parts of respective H2 HORs or similar sequences. Although the common origin of some of their monomers is quite clear, as in the trees they form branches with considerable common stem, they are more likely to be different re-amplifications of the same divergent material, which was different on each chromosome, possibly different local amplifications of S1C10/12H1. To substantiate the latter scenario, detailed study of all S1C10/12H1 arrays in different chromosomal locations focused on their chromosome specificity is needed. It should also be noted that the more divergent or rearranged copies of modern SF1 HORs located for instance on the array edges may tend to class as progenitor S1C10/12H1. So, some of the very small pieces of the latter that often flank modern SF1 arrays may not appear to be genuine.

### 2.12. Overview of SF1 HOR data

SF1 is represented by 7 different live HORs, of which one is located on 3 chromosomes (S1C1/5/19H1L), and another (S1C10H1L) has 2 related dead sequence variants (S1C10H1-B and -C). So, the total of live HORs and their close relatives is 9. All of them are modern except for S1C6H1L and a piece of 3 archaic monomers (monomers 8, 9 and 10) in S1C3H1L. Additionally, 4 homogenous pseudocentromeric HORs that are not obviously and directly related to live HORs are located in chromosomes 3, 5, 10 and 12 (S1C3H2, S1C5H2, S1C10H2 and S1C12H2). Of these, S1C3H2 is archaic. Besides, there are 4 different divergent HORs, 2 of which are located on single chromosomes (S1C3H3 and S1C12H3), one is found mainly on 2 pairs of chromosomes (S1C10/12H1) with short pieces in few other chromosomes, and one is located on many chromosomes (S1CMH1; chromosomes 3, 6, 7, 8, 10, 12, and 20). Of divergent HORs only S1C10/12H1d is modern, others are archaic. Note that the arrays of minor HORs are located in the same chromosomes which have live SF1 HORs with the exception of chromosomes 8 and 20. Altogether, there are 17 SF1 HORs of which 5 are archaic and one has a piece of archaic sequence. The detailed data on the HOR structure are shown in Fig. S1 where the obviously similar monomers within a HOR are shown by color in the first tree and in a special color panel. More loose relationships between the monomers of the different HORs are summarized in Fig. S2a and the comments on specific HORs are provided in Supplementary note 1.

### 2.13. Technical description of annotation pipeline

Below we provide description of each technical step which we used to build HumAS-HMMER and to generate the HOR annotation of hg38 genome assembly and related statistics. The steps are summarized in Fig. S1.

#### 2.13.1. Building HumAS-HMMER HOR module

1. *RM analysis and extraction of a sample region.* We self-aligned each RM and inspected the dotplot in Gepard v1.4 [22] under default parameters in order to visualize HORs. We then extracted a sample region containing three to five copies of a homogeneous HOR (which was supposed to be present in each RM) so that the most typical of these often slightly different copies could later be chosen as the “master HOR” copy (see below). In cases of divergent HORs, where, given small sample and low identity, it was difficult to understand the composition of the basic HOR, we analysed the whole RM or one entire genomic region (clone or contig) with as many copies as was possible.
2. *Processing of a sample region.* In order to split the sequence into AS monomers we had to convert FASTA to Genbank format with HMMER-Easel/SQUID tool “sreformat” (http://eddylab.org), translate newlines to DOS/Windows and feed it to the PERCON program v4.35.1sR2 [1] using the option which allowed to print out separate monomers. Next we extracted monomers from PERCON output, filtered by length and for duplicate names to remove duplicate monomers that were formed in the overlapping areas of the 5 kb windows in which PERCON operates.
3. *Phylogenetic analysis of monomers from a sample region and selection of a master HOR.* That included alignment of monomers, construction of a tree, identification of monomer groups (branches), marking of monomers with group names, establishment of a number of monomers in a HOR and their basic order. These operations were performed as follows. Multiple sequence alignments were generated with MEGA-CC software v7.0.18 [23] using the MUSCLE algorithm [24]. To reconstruct the phylogeny, the Minimum Evolution method implemented in MEGA-CC was applied (additional settings: Model/Method=p-distance, Gaps/Missing Data Treatment=Partial deletion, Site Coverage Cutoff (%)=95, ME Heuristic Method=Close-Neighbor-Interchange (CNI)). The unrooted phylogenetic tree was saved into Newick format. MEGA Alignment Explorer [25] was used in cases where manual refinement was required. MEGA Tree Explorer was used to assign names to groups of nodes that would later become numbers of monomers within a HOR. Group names were exported to text file and viewed as a linear map, if a correction in naming was required, the process was repeated. When the basic order of the monomers was clear, one of the typical repeats was selected as a master HOR and the names of the monomers were assigned. Monomer name included the HOR name and the monomer number (e.g. S1C3H1L.1 for monomer 1 and S1C3H1L.2 for monomer 2).
4. *Preparation of HMM profiles.* From resulting FASTA alignment of the master HOR monomers HMM profiles were prepared by the method previously described [26] with modifications (two identical monomers in a profile, but only one strand is used) and formatted using hmmbuild program from the HMMER v3.0 package [27]. Every monomer of a homogenous HOR made up a single profile. For divergent HORs, a group of divergent copies of the same monomer made up a single profile (MSA profiles). In addition to profiles for SF1 HOR monomers, HMM profiles for AS SF-specific classes of monomers were added as a MSA (12 profiles total, described in [1]).
5. *Final check for identical monomers.* When all RMs were processed in the above way, the whole pool of SF1 HOR monomers used for the profiles was subjected to the final phylogenetic analysis to make sure that no HORs were identical or contained identical monomers. If indistinguishable monomers were found in different HORs they would have to be marked, but so far no such cases were observed.
6. *Initial version of HumAS-HMMER is ready to use.* After the above steps the initial version of the tool was applied to analyse the assembly or its parts with the nhmmer tool of the 1.HMMER v3.1b2 package [28] and the output was used to make the HOR-track.

### 2.13.2. Building and use of a HOR-track

#### 1. Description of HMMER output

HMMER target hits table consists of one line for each different query/target comparison (where target is an AS monomer in the assembly and query is a profile in HMMER) that meets the reporting thresholds. The lines are ranked by decreasing statistical significance (increasing E-value). The number of lines for each AS monomer in the assembly would equal the number of profiles in HMMER, as a monomer is compared to every profile. Each line consists of 15 space-delimited fields followed by an optional free text target sequence description. From this output we select the following: ‘target name’ ‘query name’, ‘alifrom’ and ‘ali to’ which are the start and the end of a hit in target sequence, ‘strand’ which states if the hit was found in the “+” or the “-” strand and ‘score’ with the score (in bits) for this hit, which includes the biased-composition correction.

#### 2. Conversion of the output to BED file for the HOR-track

Selected fields from the tabular output are converted to the BED format file using the script (which selects the required fields, color-codes HORs and also applies the threshold ratio of score to monomer length. In order to reduce background of non-specific hits over non-AS and non-SF1 sequences, only the scores for which this ratio was more than 0.7 were used further. If a target does not have any scores which meet the threshold, it is dropped from the list and appears in the track as not covered. This was followed by sorting of all target sequences in the file according to their coordinates in a chromosome and to the succession of chromosomes in the assembly, so that the target monomers (each with its multiple hit scores) would be arranged in the order they have in the assembly. Finally, the hits were filtered in such a way that of all overlapping hits with a given target only the hit with the best score was retained. This was done using bedmap program from the BEDOPS suite [29] (with settings: --max-element --fraction-either 0.1). In the end, each line for a target-query comparison had the name of the winning query with the highest score instead of the query originally used in comparison. Of these multiple identical lines a single line with the description of the best hit for this target monomer was picked up for the final BED file. This set of tasks was implemented via a script to run on SLURM Cluster [30] (https://github.com/enigene/HumAS-HMMER).

#### 3. Viewing the track

The track is self-explanatory, as in the full-view mode each monomer is named according to the rules of our new HOR naming system (see section 2.3 above) and monomers of each different HOR have a different color. In the dense mode, HOR arrays appear continuously covered with respective HOR color and any mixing of colors indicates either the presence of a segment of alien HOR sequence or occasional misrecognition of a monomer by HumAS-HMMER. In the first case, there would usually be few monomers of different color in the correct order, and in the second case a single monomer of a different color. The current level of misrecognition may be improved in the future by further application of the color separation procedure described above. Note that for sequence variants of the same HOR (different hyphenated letters of the same name) significant color mixing is expected. The monomers in such HORs were named in such a way that this admixture would not create the numerical chaos and the order of monomers in a HOR could be read correctly despite mixing.

#### 4. Using the track for data mining

Resulting basic annotation (‘track’) for the human GRCh38/hg38 assembly (RefSeq accession GCF_000001405.26) was inspected in the Web service UCSC Genome Browser [31] (https://genome.ucsc.edu/) or in a local installation of the GBiB [32]. By using the Table Browser, the track data can be analyzed in text format and filtered or transformed to generate various statistics. For instance, different HOR monomers can be counted per individual chromosome or chromosomal region. Also, the overlaps with other tracks can be created and retrieved as a new track, DNA sequence, or in text format. In this paper, we used the overlaps between the HOR- and the SF-tracks. Reports were generated via SVASHOR package (https://github.com/enigene/SVASHOR) which contained information on the number of copies for each structural HOR variant in each RM, detailed representation of HOR variants (with indication of A/B type and SF class obtained from PERCON SF-track [1], and HOR monomer names obtained from the HOR-track), full map of RM, and rendering of simple trees which was done by a special script (https://github.com/enigene/print-phylo-newick-tree). In complex cases tree images were generated manually using MEGA Tree Explorer.

### 2.14. Criteria of annotation success

The annotation of SF1 AS HORs which we report here has been executed using a very simplistic approach and is essentially self explanatory. The correctness of annotation was self-evident as indicated by completeness of coverage, color separation and numerical order (as opposed to chaos) in the track which meant that monomers in basic HORs tended to go in correct numerical succession. These three features were percepted as the evidence of good coverage. Only in cases of divergent HORs which form a tiny (about 3%) proportion of SF1, reduced numerical order was often observed and there it could well be explained by degradation via deletions and other rearrangements. Also, as some divergent HORs tend to reside in more than one chromosome, the study of HOR arrays in each location is due. It may reveal multiple chromosome-specific or location-specific subfamilies within the divergent HOR families, similar to sequence variants in homogenous HORs. Then, the introduction of each separate subfamily as a set of HMM profiles could improve the color separation in divergent HOR arrays. Overall, the near complete coverage of SF1 by HOR-specific HMM profiles was a demonstration of annotation success given that a satisfactory numerical order was achieved in RMs.

The background of false hits outside SF1 arrays in the track was sufficiently low. It was located mainly in SF4+ (0.2% of total SF1 hits) where it involved thin clouds of isolated low score hits which usually did not cover a whole monomer, and in SF5 (could not be easily estimated, but presumed to be also about 0.2%) where the hits formed similar clouds, but with somewhat higher scores. The low-score false coverage in M1+ (mean score to length ratio 0.72) was an expected result of our low-threshold strategy. It could be reduced by increasing our score to length ratio threshold (currently 0.7) but only at the expense of somewhat reduced coverage in divergent SF1 HOR arrays. The higher score false coverage in SF5 (mean score to length ratio could not be determined, in selected hits studied it was 0.8 - 0.9) could be eliminated only by a dramatic threshold increase which would be even more costly. Our preliminary data showed that the false coverage was mainly a problem of inadequate competing class-specific profiles for SF5 and SF4+. In fact, upon introduction of separate profiles for each colored layer identified within SF4+ [2], which provided a tighter fit for the M1+ monomers, the false coverage of SF4+ was greatly reduced. These profiles will be reported separately.

Annotation of SF1 HORs performed in this work was done with a very low threshold and every SF1 monomer had identified with a certain single-monomer or MSA HMM profile, to which it had the best match. In the final HOR-track version, no monomers which identified with J1 or J2 in PERCON SF-track remained not covered by SF1 HOR profiles. Thus, effectively, in the final version, the class-specific profiles played no role and could have been switched off. If we used a higher threshold and filtered for high-score similarity, only the homogenous HORs would get identified (mean score to length ratio 1.15). With no high-score filtering we were also able to identify divergent HOR arrays where the similarity scores were significantly lower (mean score to length ratio 0.97). However, the color separation and coverage in divergent arrays expectedly were not as good as in homogenous HOR arrays.

As a whole, our annotation have visualized the known and expected structure of AS arrays, including the large homogenous cores of the live centromeres, and a number of additional smaller homogeneous arrays percepted as pseudocentromeres. Perhaps in some (or many) cases the latter could be half-dead centromeres, which would function as centromeric epialleles [4, 5] in a small part of individuals within a population. However, two new important partitions were amply revealed as a result of our annotation of SF1, which were archaic versus modern HORs and homogenous versus divergent HORs. Our comments on both are presented in Supplementary Note 2. The latter also contains the comments on detection of polymorphic variants of AS HORs using HumAS-HMMER.

### 2.15. Perspective

In this paper we presented the first installment of the HMMER-based AS HOR annotation of the latest hg38 human genome assembly, which covered AS SF1 arrays. The principal features of this annotation were the monomer-by-monomer processing of the HORs and a very low identity threshold used. These features have allowed to start addressing such previously unidentified problems as hybrid monomers and divergent HORs. The slight setback noted which would require further work was some false recognition background in non-SF1 arrays. We surmised that the manner of annotation we have chosen worked fine for the human genome assembly and could clearly be further improved and developed in the future. Similar annotation of the other AS SFs has been largely accomplished and will be published shortly. We believe the HumAS-HMMER HOR module can also be used for annotation of the short deep sequencing reads, but should first be carefully evaluated for the effect of using short monomer fragments instead of the whole monomers used in this work.

### 2.16. Supplementary sequence files attached to this paper

We attached to this paper a number of files with aligned monomers which can be useful for different purposes, so that the trees or versions of HumAS-HMMER could be constructed as needed and the conclusions of this paper verified. These and additional files can also be viewed at our repository (https://github.com/enigene/Files-and-scripts-used-in-SF1-HORs-in-hg38-article). Files attached to this paper are described below.

#### 2.16.1. HMM profiles file

HMM profiles prepared by us and used in HumAS-HMMER. A single file ready to be installed in HMMER. 581 profiles including 12 class profiles for SFs other than SF1 which are described in [1].

#### 2.16.2. Alignment file 1

Aligned monomers as they were used for the profiles (HOR-specific alignment). The monomers of each HOR were aligned together, but separately for different HORs, and then pooled. So, the monomers of different HORs may have different length and some characteristic deletions (such as in positions 52 and 68 of J2 monomers) may be located variously. Provides unchanged sequence without any alignment deletions for every monomer as a reference. The coordinates of each monomer in the assembly can thus be obtained by finding 100% identical monomer using BLAT search in UCSC Browser. Note that we used single-monomer profiles for homogeneous HORs and MSA profiles for divergent HORs (as described in section 2.2). Monomers from this file were used to construct the trees shown in Fig. S1 (the first and the second tree for each HOR).

#### 2.16.3. Alignment file 2

All the monomers used in this paper aligned together and adjusted to the length of 171 bp (i.e. some deletions may have been made). Characteristic deletions (such as in positions 52 and 68 of J2 monomers) are located uniformly. Sorted in A and B types. The monomers are aligned together with addition of SF class consensus and type consensus monomers in latest edition and the J1m and J2m consensus monomers derived in this paper.

#### 2.16.4. Alignment file 3

Only the monomers of master HORs for homogenous HORs and only consensus monomers for divergent HORs. Sorted in A and B types. Additional consensus monomers were included as in file 2. Monomers from this set were used for the trees shown in Fig. S2. For the haplotype analysis shown in Table S1 the same monomers were used with modifications to consensus sequences listed in Table S3.

## Correspondence

IAA

EIR

## Acknowledgements

Y.B. Yurov has deceased when this work was already concluded. With sadness and gratitude the remaining authors acknowledge his memory.

## Supplementary material

- Fig. S1. The structure of SF1 HORs.
- Fig. S2. Relationships between the monomers of different SF1 HORs.
- Fig. S3. Schematic representation of HumAS-HMMER SF1 HOR module annotation pipeline.
- Tables S1 - S4.
- Supplementary Notes 1 and 2, description of Supplementary Figures and Tables.
- HMM profiles file
- Alignment file 1
- Alignment file 2
- Alignment file 3

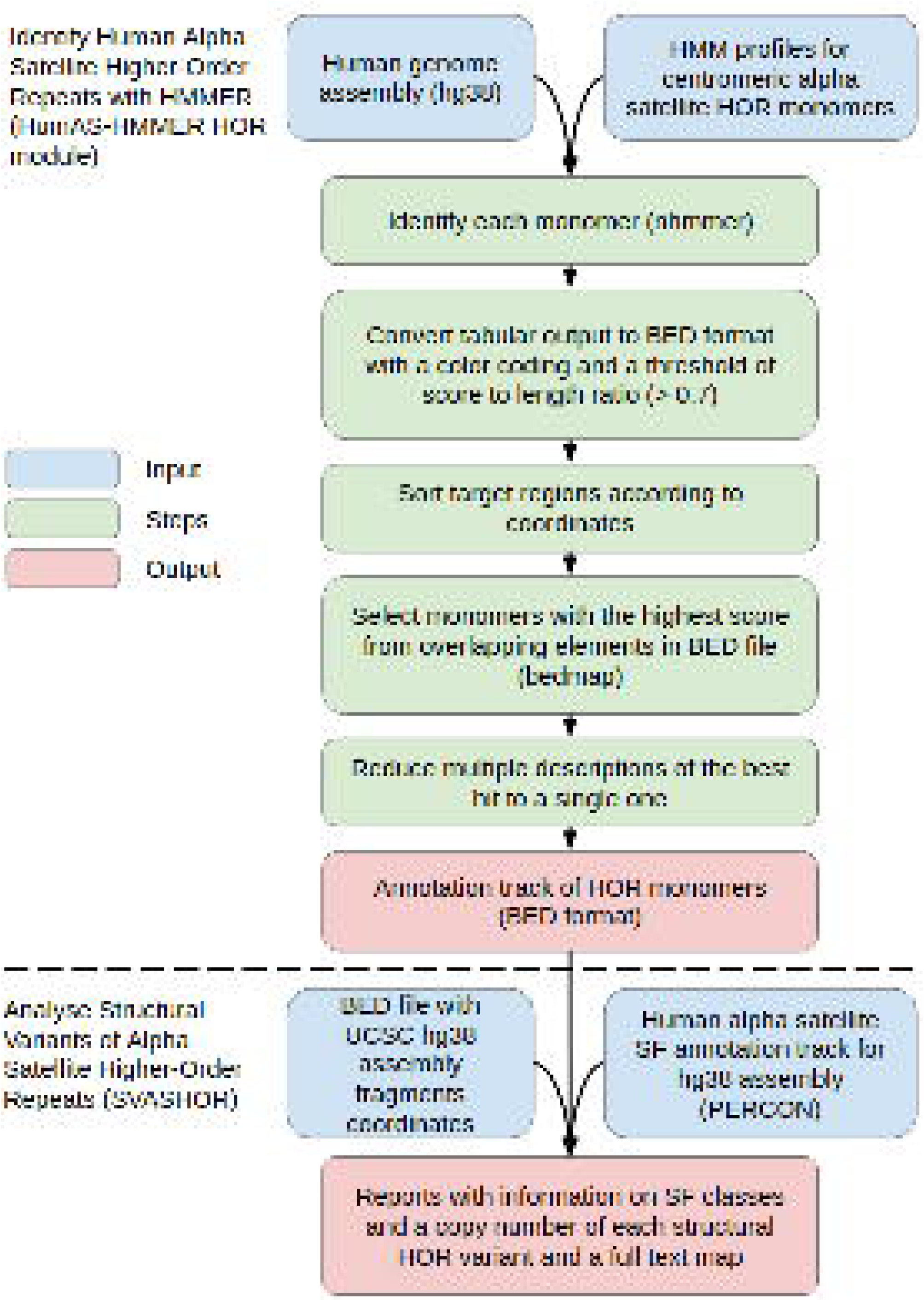

